# *Quantum pBac*: An effective, high-capacity *piggyBac*-based gene integration vector system for unlocking gene therapy potential

**DOI:** 10.1101/2022.04.29.490002

**Authors:** Wei-Kai Hua, Jeff C. Hsu, Yi-Chun Chen, Peter S. Chang, Kuo-Lan Karen Wen, Po-Nan Wang, Yi-Shan Yu, Ying-Chun Chen, I-Cheng Cheng, Sareina Chiung-Yuan Wu

## Abstract

Recent advances in gene therapy have brought novel treatment options for cancer. However, the full potential of this approach has yet to be unlocked due to the safety concerns and limited payload capacity of commonly utilized viral vectors. Virus-free DNA transposons, including *piggyBac*, have potential to obviate these shortcomings. In this study, we improved a previously developed modified *piggyBac* system with superior transposition efficiency. We demonstrated that the internal domain sequences (IDS) within the 3’ terminal repeat domain of hyperactive *piggyBac* (*hyPB*) donor vector contain dominant enhancer elements. Plasmid-free donor vector devoid of IDS was used in conjunction with a helper plasmid expressing *Quantum PBase*™ v2 to generate an optimal *piggyBac* system, *Quantum pBac*™ (*qPB*), for use in T cells. Cells transfected with *qPB* expressing CD20/CD19 CAR outperformed those transfected with the same donor vector and plasmid expressing *hyPB* transposase in terms of CAR-T cell production. Importantly, *qPB* yielded mainly CD8^+^ CAR-T_SCM_ cells, and the *qPB-* induced CAR-T cells effectively eliminated CD20/CD19-expressing tumor cells both *in vitro* and *in vivo*. Our findings confirm *qPB* as a promising virus-free vector system with a payload capacity to incorporate multiple genes. This system is highly efficient and potentially safe for mediating transgene integration.

## Introduction

Gene therapy often requires stable, long-term expression of therapeutic transgene(s) in cells. Such prolonged expression can be accomplished by engineering cells either *ex vivo* or *in vivo* using viral or non-viral vector systems. Viral vectors are most commonly used for gene therapy due to their high efficiencies of gene delivery and integration, which confer stable and long-term gene expression. However, viral vectors have several intrinsic limitations, including: (1) limited payload capacity that severely restricts the repertoire of genes that can be integrated^1^; (2) genotoxicity arising from preferential integration into sites near or within active gene loci that may negatively impact the expression and/or function(s) of genes^2–7^; (3) a cellular proclivity for silencing genes introduced by viral vectors, presumably due to cellular immunity^8,9^; and (4) safety concerns related to immunogenicity of viral vectors^10^. Additionally, production of viral vectors for clinical trials is costly, time-consuming (> 6 months), and supply-constrained, presenting significant obstacles to the routine use of viral vectors in medical practice^11^.

In recent years, many advances have been made in non-viral gene delivery technologies, such as virus-free DNA transposons. These advancements may be capable of bypassing the shortcomings of viral vectors, and transposon technology has emerged as an especially promising vector system for gene therapy^12,13^. The great promise of transposon technology is largely due to its effective gene integration capability^14^. DNA transposons, also known as mobile elements or jumping genes, are genetic elements with the ability to transverse the genome through a “cut-and-paste” mechanism. In nature, a simple DNA transposon contains a transposase gene flanked by terminal repeat sequences. During the transposition process, the ability of transposase to act on virtually any DNA sequence flanked by the terminal repeat sequences makes DNA transposons particularly attractive as vectors for gene therapy. To turn DNA transposons into tools for genetic engineering, researchers have developed controllable bi-component vector systems, consisting of (1) a helper plasmid expressing the transposase and (2) a donor plasmid with exogenous DNA of interest flanked by the transposon terminal repeat sequences. Currently, *Sleeping Beauty* and *piggyBac* are thought to be the most promising DNA transposons for human gene therapy and have been clinically explored as vectors for several CAR-T cell therapies. Unlike *Sleeping Beauty*, which was reconstructed from the salmon genome^15,16^, *piggyBac* was derived from the cabbage looper moth *Trichoplusia ni* and is naturally active in humans^17–19^. By introducing amino acid mutations to the transposase, a hyperactive *piggyBac* (*hyPB*) transposase, (*hyPBase*) and two hyperactive transposases of *Sleeping Beauty* (SB100X and hyperactive SB100X, which is 30% more active than SB100X) were developed^20–24^. When an exogenous gene is delivered to primary human T cells *ex vivo, hyPBase* can increase transposition efficiency by 2-to 3-fold compared with *piggyBac* or SB100X transposases. Previously, we shortened the *piggyBac* terminal repeat domain (TRD) sequences and observed a 2.6-fold increase in transposition activity mediated by *piggyBac* in HEK293 cells^25^. We also demonstrated that the activity of *hyPBase* can be increased by another 2-to 3-fold by fusing the transposase with various peptides^26^. In this study, we address whether the *piggyBac* system can be further developed for therapeutic application, including increasing its cargo capacity relative to those of currently available *piggyBac* systems. To do so, we generated *Quantum pBac*™ (*qPB*), a binary *piggyBac* system comprising a plasmid-free donor vector and a helper plasmid expressing the molecular-engineered *hyPBase, Quantum PBase™* (*qPBase*) *v2*. This simple, robust and potentially safe vector system can be used to generate potent CD19/CD20 dual-targeting CAR-T cells for treatment of B cell malignancies.

## Results

### Micro-*piggyBac* possesses significantly lower enhancer activity compared to mini-*piggyBac*

In the initial clinical trial of retrovirus-based gene therapy for SCID-X1, there was evidence of malignancies caused by vector-mediated insertional activation of proto-oncogenes. The currently available minimal *piggyBac* transposon vector, designated as mini-*piggyBac* here, contains 5’ (244 bp) and 3’ (313 bp) TRD *cis* elements, which are transposed along with gene of interest into the genome. Each TRD contains a Terminal Inverted Repeat (TIR; also known as inverted minimal terminal repeats) sequence and an internal domain sequence (IDS)^27^. To minimize the potential risk of insertional mutagenesis caused by gene delivery vectors in gene therapy, we had previously generated micro*-piggyBac*. This vector contains the 5’ (67 bp) and 3’ (40 bp) TIR sequences of mini-*piggyBac*, but the respective

IDS are absent (Figure 1)^25^. Nevertheless, it remained unclear whether the TRDs of mini-*piggyBac* and/or TIRs of micro-*piggyBac* harbor enhancer and/or silencer activities. To address this issue, a panel of luciferase reporter constructs containing individual TRD or TIR sequences was generated and examined in insect Sf9 cells, human HEK293 cells, and human Jurkat T cells (Figure 1).

**Figure 1.**
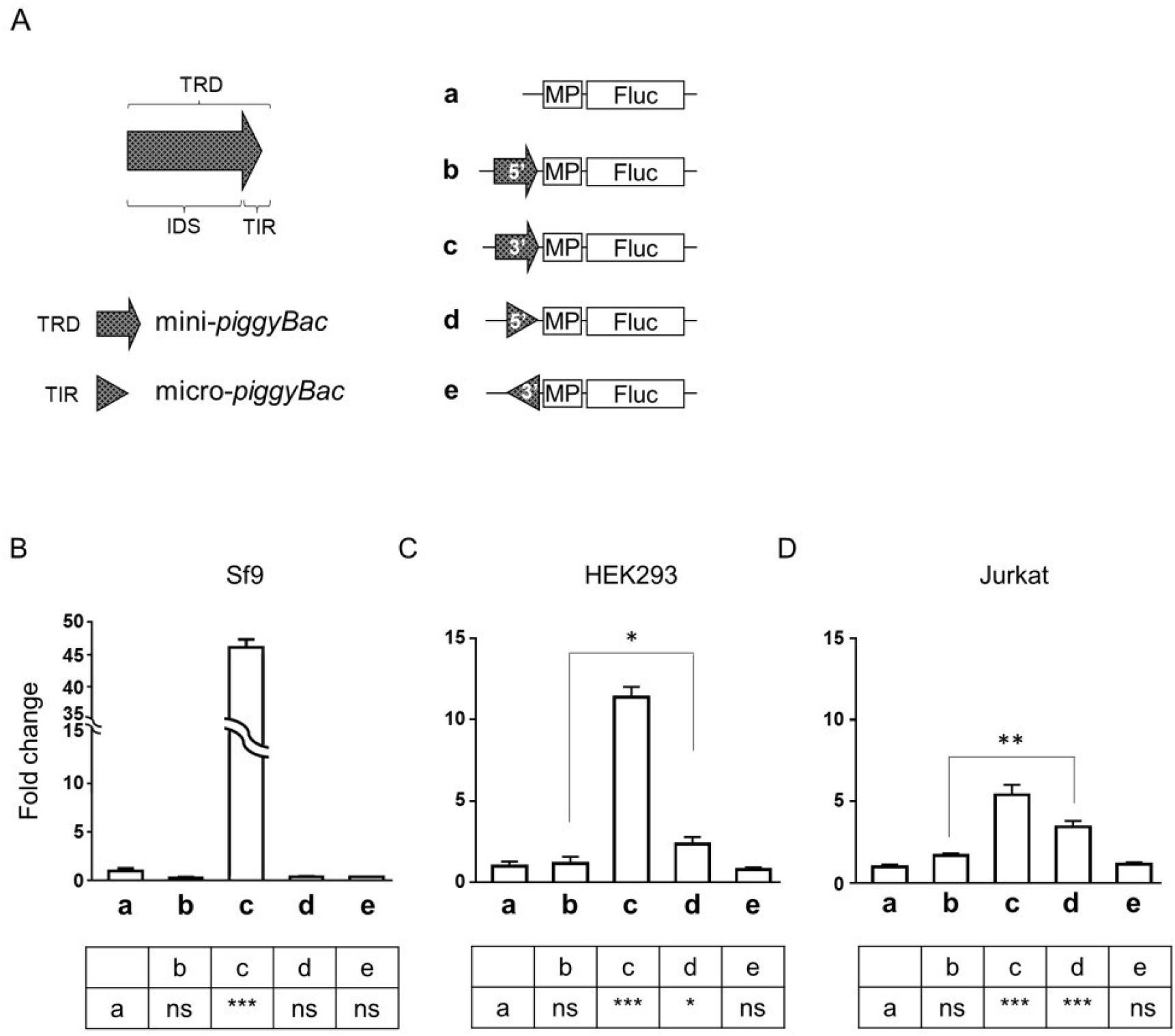
Enhancer/silencer activity of mini-*piggyBac* TRDs and micro-*piggyBac* TIRs. (A) Schematic depicts the panel of luciferase reporter constructs containing a CMV minimal promoter (MP) and firefly luciferase (*Fluc*) gene, with or without the 5’ or 3’ TRDs from mini-*piggyBac* or 5’ or 3’ TIRs from micro-*piggyBac* inserted upstream of MP. Luciferase activities exhibited by the reporter constructs of (A) in (B) Sf9, (C) HEK293, and (D) Jurkat cells. Results are shown as mean fold changes in luciferase activity ± standard deviation (SD; normalized first to Renilla luciferase activity and then to the luciferase activity obtained from cells transfected with construct a). Statistical analysis comparing luciferase activities obtained from cells transfected with construct **a** and those from cells transfected with constructs **b**-**e** are summarized in the lower panels (boxed regions). * p < 0.05, ** p < 0.01, *** p < 0.001. N = 3 (triplicate).

Compared to the control (Figure 1A, construct **a**), the 3’ TRD of mini-*piggyBac* (construct **c**) but not 3’ TIR of micro-*piggyBac* (construct **e**) produced significantly higher luciferase activity across all three cell types (Figure 1B-1D), suggesting that enhancer activity is present in the 3’ IDS of mini-*piggyBac*. On the other hand, a slightly yet significantly enhanced luciferase activity was detected in 5’ TIR of micro-*piggyBac* but not 5’ TRD of mini-*piggyBac* in both HEK293 and Jurkat cells. This result suggests the presence of a minimal level of enhanced activity in the 5’ TIR and/or silencer activity in the 5’ IDS of TRD (Figure 1B-1D). Taken together, the data suggest that micro-*piggyBac* possesses significantly lower enhancer activity compared to mini-*piggyBac*.

### Shortening of donor vector backbone in combination with *Quantum PBase™* (*qPBase*) *v2* enhances the transposition activity of micro-*piggyBac* in T cells

The significantly reduced enhancer activity of TIRs suggested that micro-*piggyBac* may be safer than mini-*piggyBac* for gene therapy applications. We therefore focused on micro-*piggyBac* and determined whether its transposition activity may be further enhanced by shortening the donor vector backbone (i.e., the sequences outside the TIR-spanning region). We constructed two donor vectors named micro-*piggyBac*-Short and micro-*piggyBac*-Long. Both of these vectors contain TIRs (micro) and either retains (Long) or is devoid of (Short) replication components in its backbone, namely the replication origin and antibiotic genes. A third donor vector, mini-*piggyBac* Long, which contains TRDs (mini) and retains the replication components in its backbone (Long), was also constructed for comparison; its combination with helper plasmid expressing *hyPBase* (collectively called “hyperactive *piggyBac*” or “*hyPB*”) is currently the most advanced *piggyBac* system available (Figure 2A). We co-electroporated HEK293, Jurkat or human primary T cells with different combinations of these donor vectors and helper plasmids expressing wild-type *PBase* (pCMV-Myc-PBase), *hyPBase* (pCMV-HA-hyPBase), and *qPBase v1* or *v2* (pCMV-qPBase_v1 and pCMV-qPBase_v2, respectively). We then determined and compared the transposition efficiencies of these donor vector and helper plasmid combinations (Figure 2A).

**Figure 2.**
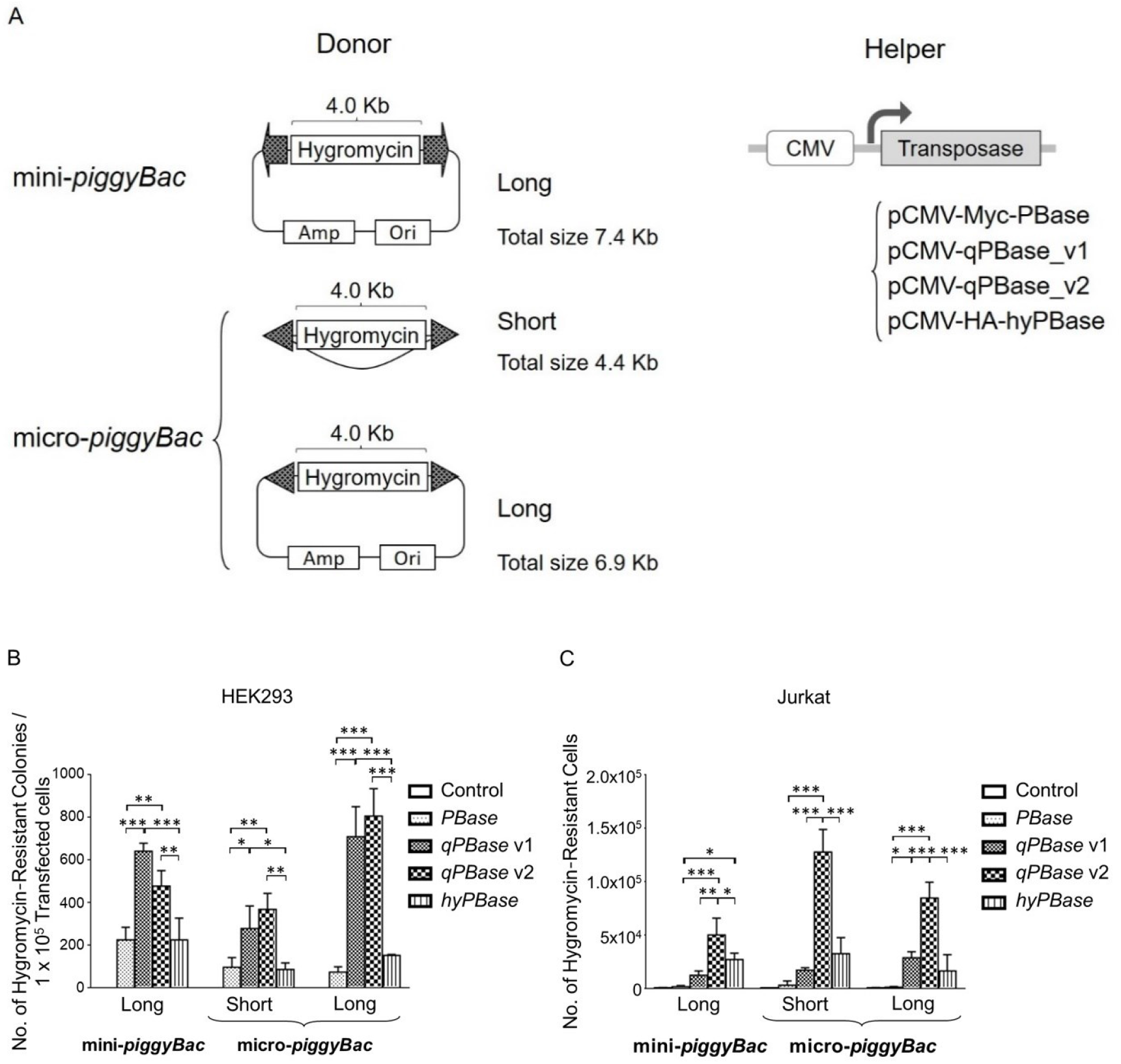
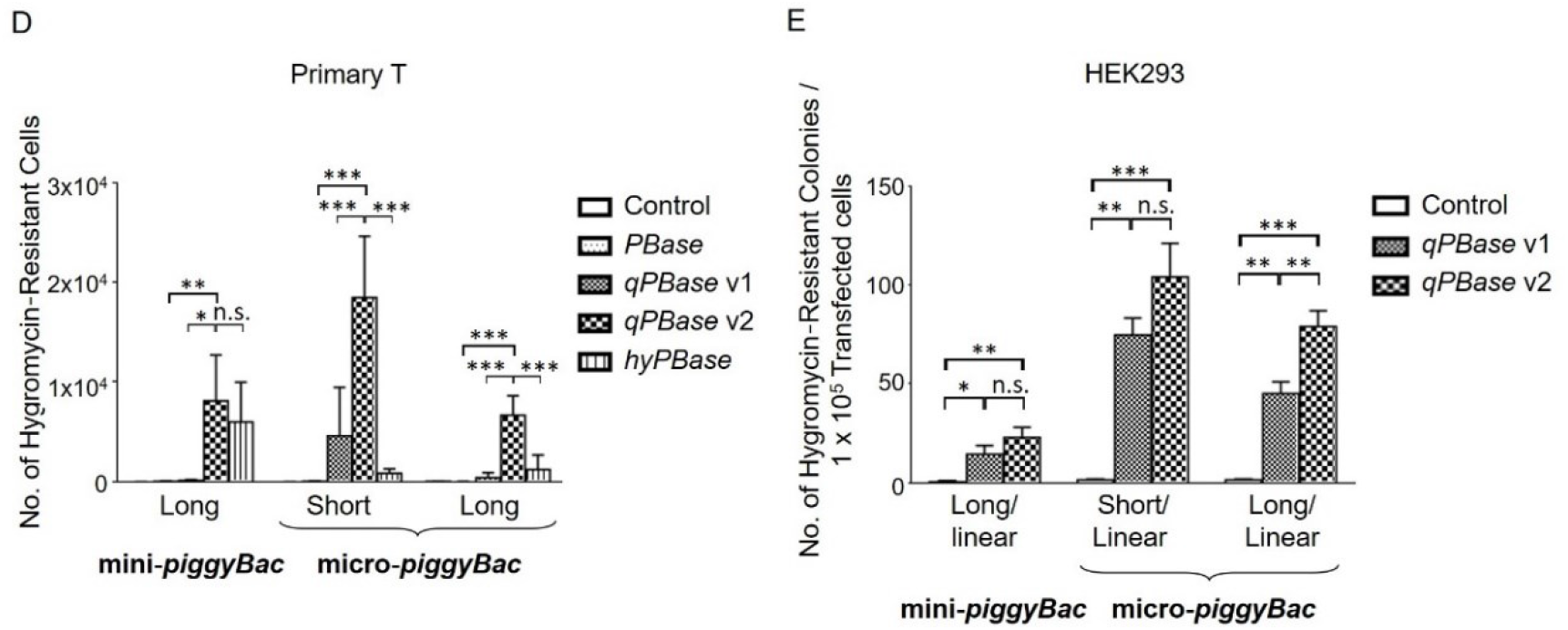
Transposition activities of various combinations of donor vectors and helper plasmids in human cells. (A) Schematic depicts the *piggyBac* system with various donor vectors and helper plasmids. Transposition activity of the indicated combination of donor vector and helper plasmid in (B) HEK293, (C) Jurkat, and (D) primary T cells. (E) Transposition activity of the indicated combination of linearized donor vector and helper plasmid in HEK293 cells. Results are shown as mean ± SD (B-E). n.s. not statistically significant, * p < 0.05, ** p < 0.01, *** p < 0.001. N = 3 (triplicate).

As shown, *hyPBase*, either in combination with mini-*piggyBac*-Long or micro-*piggyBac*-Short or -Long, markedly enhanced transposition compared to *PBase* in T cells but not in HEK293 cells. On the other hand, *qPBase* v1 markedly enhanced transposition compared to *hyPBase* in HEK293 cells but not in T cells (Figure 2B-2D). These results suggest that the transposition activity of *hyPBase* and *qPBase v1* are likely cell-type-dependent. We also found that *qPBase v2* mediated the highest transposition activity in nearly all of the tested combinations and cell types. Importantly, when *qPBase v2* is accompanied by micro-*piggyBac*-Short donor vector, its transposition activity is by far the highest in both Jurkat and primary T cells, and this combination is clearly superior to the hyperactive *piggyBac* system, *hyPBase* in combination with mini-*piggyBac*-Long (Figures 2C and 2D).

### Micro-*piggyBac* is superior to mini-*piggyBac* for advancing adeno-*piggyBac* hybrid vector for gene therapy

Even though *piggyBac* is capable of integrating sizable DNA (> 100 kb), its genome engineering efficiency is restricted by the effectiveness of gene delivery methods. Electroporation is a particularly effective virus-free gene delivery method commonly used in gene therapy. However, its transfection efficiency is inversely correlated with the size of DNA delivered, as cell damage may be caused by introduction of larger transgenes. Thus, to alleviate electroporation-associated restrictions imposed on the *piggyBac* system, an adenovirus-*piggyBac* hybrid system (Ad-iPB7) was developed^28^. Since the adenovirus genome exists in a linear form, we examined the transposition activity of linearized forms of mini-*piggyBac*-Long, micro-*piggyBac*-Short and micro-*piggyBac*-Long along with either *qPBase v1* or *qPBase v2*. As shown in Figure 2E, both micro-*piggyBac*-Short and micro-*piggyBac*-Long were superior to mini-*piggyBac*-Long. Moreover, when linearized micro-*piggyBac*-Long donor vector was combined with *qPBase v2*, a significantly greater transposition efficiency was observed relative to that when *qPBase v1* transposase was used (Figure 2E).

### Minicircle micro-*piggyBac* is significantly more active than its parental counterpart in both Jurkat and primary T cells

A minicircle DNA is a plasmid-free nucleic acid sequence that is normally generated by an *in vivo* recombination process, which excises prokaryotic plasmid components such as the antibiotic resistance gene and bacterial origin of replication. Minicircle DNA constructs offer several advantages, including enhancements in gene delivery efficiency and stable transgene expression. These advantages may be attributed to the small vector size of minicircle DNA, the absence of antibiotic resistance genes, and the lack of gene silencing induced by bacterial backbone sequences required for plasmid replication. We demonstrated that removal of backbone sequences outside the TIR-spanning regions via a cloning procedure (which shortens the vector length) enhanced transposition efficiency (Figure 2). Therefore, we next used the Mark Kay minicircle system to generate a minicircle form of the donor vector (Q-tdTomato-IRES-hygro) from its parental plasmid counterpart (pQ-tdTomato-IRES-hygro) (Figure 3A)^29^. We carried out the same transposition assay as described for Figure 2 and determined whether transposition of donor vector in the minicircle form into the T cell genome would be more efficient than that of its parental plasmid counterpart.

**Figure 3.**
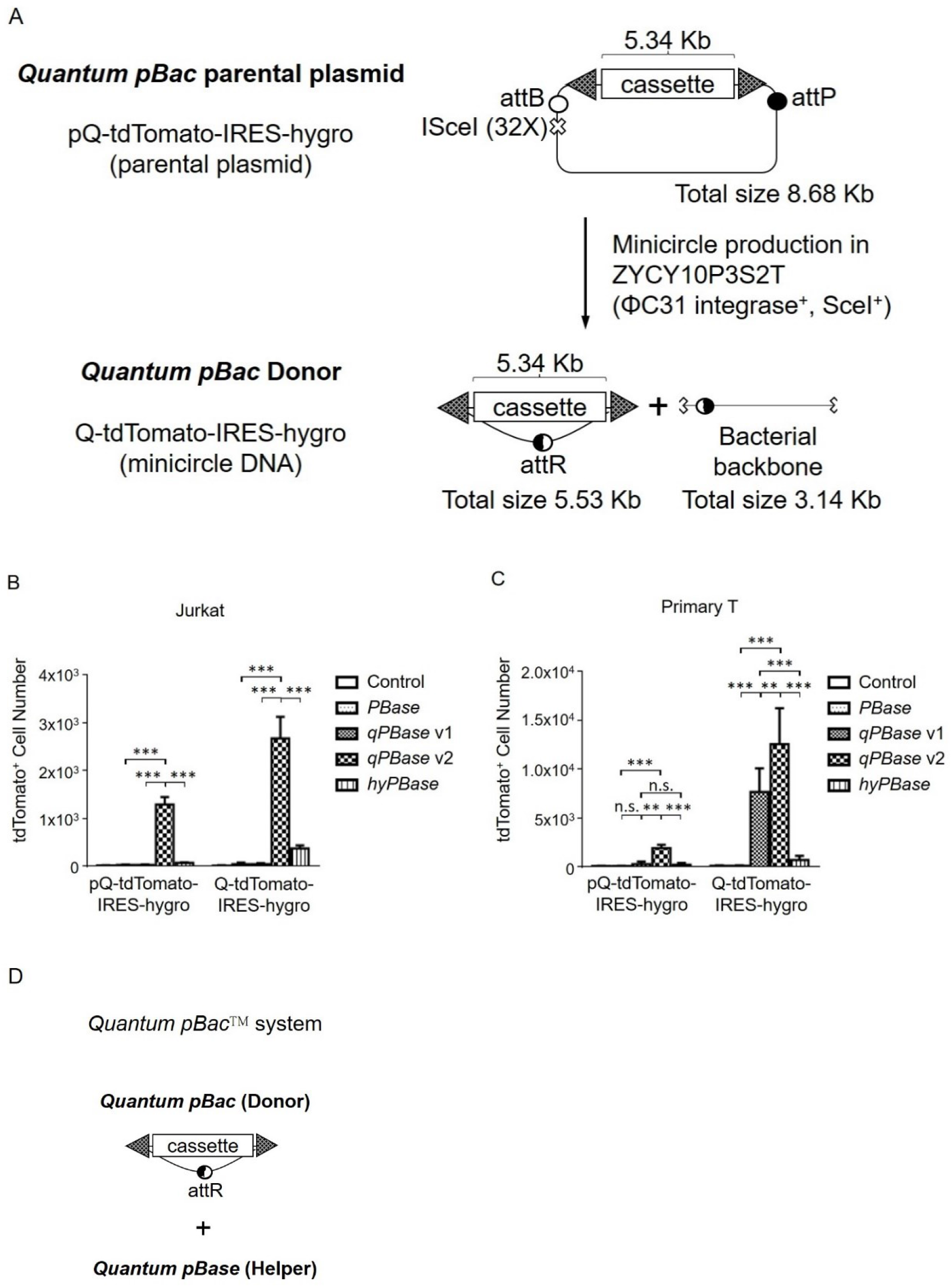
Transposition activity of various transposases with donor vector in parental plasmid or minicircle counterpart forms. (A) Schematic depicts the generation of the *Quantum pBac*™ (*qPB*) donor vector. A gene cassette carries *tdTomato* and *hygromycin* genes linked by IRES (Q-tdTomato-IRES-hygro) from the parental plasmid (pQ-tdTomato-IRES-hygro). Transposition activities of the indicated combinations of pQ-tdTomato-IRES-hygro or Q-tdTomato-IRES-hygro donor vectors and helper plasmids in (B) Jurkat and (C) primary T cells. (D) Schematic depicts the two-component *qPB* system. Results are shown as mean ± SD. n.s. not statistically significant, ** p < 0.01, *** p < 0.001. N = 3 (triplicate).

When donor vectors of parental plasmid (pQ-tdTomato-IRES-hygro) and minicircle (Q-tdTomato-IRES-hygro) forms were compared, it was clear that minicircle donor vector mediated higher transposition efficiencies than the parental plasmid form, irrespective of which helper plasmid was co-electroporated. Moreover, cells co-electroporated with minicircle donor vector and helper plasmid expressing *qPBase v2* consistently exhibited significantly higher transposition activity than those receiving all other combinations, including those with the helper plasmid expressing *hyPBase*. Based on these data in both Jurkat (Figure 3B) and primary T cells (Figure 3C), we selected micro-*piggyBac* vector in the minicircle DNA form as the donor vector and a series of recombinant *qPBase* (including *qPBase v1* and *v2*) as the helper plasmids to form the *Quantum pBac*™ (*qPB*) system (Figure 3D).

### When combined with CD20/CD19 CAR *Quantum pBac*™ (*qPB*) donor vector, *Quantum PBase*™ (*qPBase v2*) outperforms *hyPBase* in CAR-T production

We next evaluated the performance of CAR-T cells produced by electroporating activated human primary T cells with vectors of the *qPB* system.

To do so, we used anti-CD20/CD19 CAR as the transgene cassette of *qPB* donor vector and helper plasmids that encode either *hyPBase* or *qPBase v2*. (Figure 4A).

**Figure 4.**
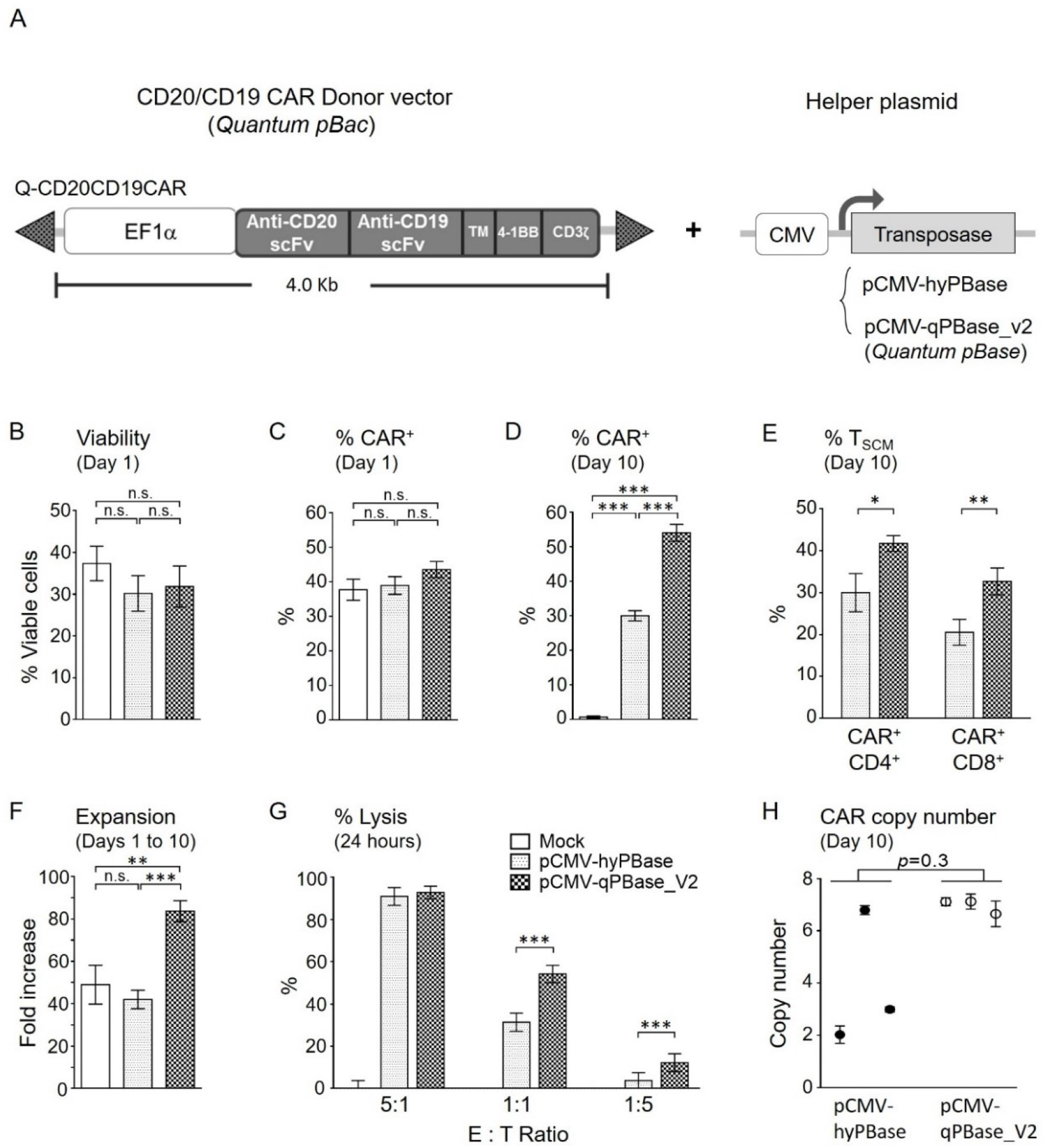
CAR-T cell production using *Quantum pBac*™ (*qPB*) donor vector and helper plasmid. (A) Schematic depicts the CD20/CD19 CAR *qPB* donor vector and helper plasmid pCMV-hyPBase or pCMV-qPBase_v2. Characterization of (B) cell viabilities one day after electroporation (Day 1). Percentages of CAR^+^ cells on (C) day 1 and (D) day 10 following electroporation of activated primary T cells. (E) The percentages of CD4 and CD8 CAR^+^ T_SCM_ cells on day 10. (F) Fold expansion of cells after 10 days of culture, (G) cytotoxicity of day 10 CAR-T cells on Raji target cells at the indicated E:T ratios, and (H) CAR copy number after 10 days of culture. Results are shown as mean ± SD (B-H). n.s., p=0.3 not statistically significant, * p < 0.05, ** p < 0.01, *** p < 0.001. N = 3 (triplicate) except in (H) where testing of each sample was repeated three times per sample.

One day after electroporation, no significant differences in viability or % CAR^+^ were found among cells electroporated with CD20/CD19 CAR *qPB* donor vector and control helper plasmid (pCDNA3.1, Mock control) and those with helper plasmid expressing either *hyPBase* or *qPBase v2* (Figure 4B, 4C). Notably, after 10 days of culture, significantly more CAR^+^ cells were observed in cultures electroporated with CD20/CD19 CAR *qPB* donor vector and helper plasmid expressing *qPBase v2* compared to *hyPBase* (Figure 4D). Moreover, the expansion and cytotoxic effects of cells electroporated with CD20/CD19 CAR *qPB* donor vector and helper plasmid expressing *qPBase v2* were also significantly greater than those seen in the other two groups of cells (Figure 4F and 4G). The CAR copy numbers between *hyPBase*- and *qPBase v2-*transposed cells were not significantly different (Figure 4H), even when CAR copy number is calculated to reflect only CAR^+^ cells (data not shown). These observations suggest that cells electroporated with helper plasmid expressing *qPBase v2* were superior with regards to their transposition efficiency, expansion capacity and anti-tumor potency than cells electroporated with *hyPBase*-expressing plasmids.

### *Quantum PBase*™ produces CAR^+^ T cells that are more highly enriched with T_SCM_ subset than those produced by *hyPBase*

Since T_SCM_ enrichment is recognized as an important potential indicator of clinical efficacy, we also compared the percentage of T_SCM_ in cells electroporated with CD20/CD19 CAR *qPB* donor vector and helper plasmid expressing either *hyPBase* or *qPBase v2* (Figure 4E). We found that in both CD4 and CD8 populations, the percentages of T_SCM_ cells were higher in cells electroporated with helper plasmid expressing *qPBase v2* than *hyPBase*. Next, we focused on the *qPB* system (*qPB* donor vector with *qPBase v2* helper plasmid) and sought to determine whether there may be donor-dependent variations in CAR^+^ cell production using this system. We analyzed the performance of CAR-T cells derived from peripheral blood mononuclear cells (PBMCs) of six healthy donors (Figure 5).

**Figure 5.**
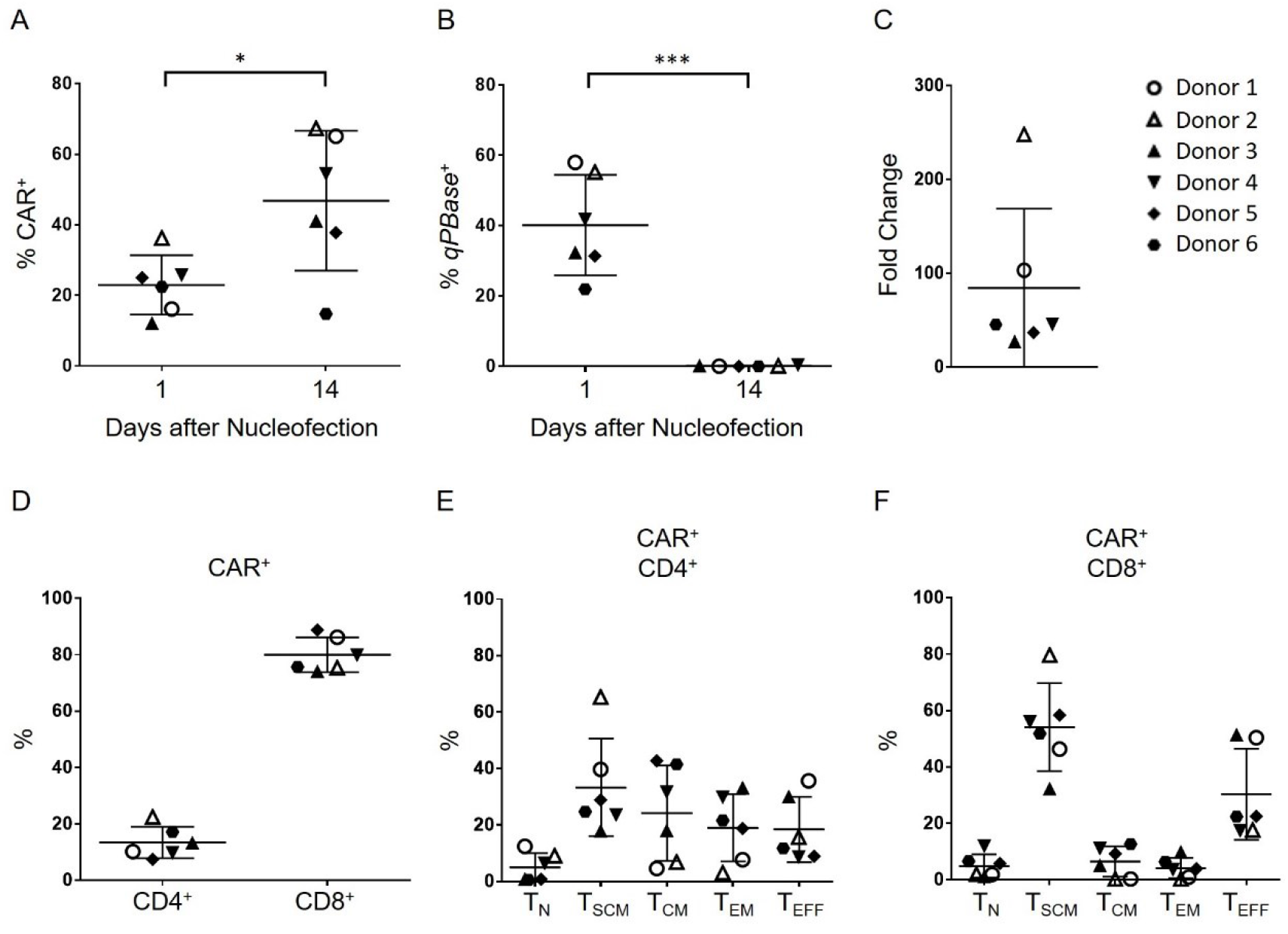
Characterization of healthy donor CAR-T cells produced using the *Quantum pBac*™ (*qPB*) system. Percentages of (A) CAR^+^ and (B) transposase^+^ (*qPBase*^+^) cells on days 1 and 14 following electroporation. (C) Fold expansion of cells and (D) distributions of CD4^+^ and CD8^+^ T cell subtypes in CAR^+^ T cells after 14 days of culture. Distributions of five T cell differentiation subsets T_N_, T_SCM_, T_CM_, T_EM_ and T_EFF_ among (E) CD4^+^ CAR^+^ and (F) CD8^+^ CAR^+^ T cells on day 14. Results were derived from cells taken from six healthy donors. Horizontal lines in (A-F) represent mean ± SD. * p < 0.05, *** p < 0.001. N = 6 PMBC donors.

As shown in Figure 5A, the average percentage of CAR^+^ T cells significantly increased from day 1 to 14 following electroporation, with only one of six donors (donor 6) exhibiting a decrease in percentage of CAR^+^ T cells. Moreover, fourteen days after electroporation, the percentage of *qPBase*^+^ cells decreased to minimal levels (< 0.4%) in all of the donor groups (Figure 5B), suggesting successful clearance of unwanted helper plasmids from T cells following completion of “cut-and-paste” gene integration. There was high variability among the PBMC donor groups in terms of the extent of CAR^+^ T cell expansion during the 14-day culture period (27-to 248-fold; Figure 5C), indicating a donor-dependent effect on expansion capacity of the cells. We also profiled the T cell subtypes (CD4^+^ and CD8^+^; Figure 5D) and T cell subsets based on differentiation stages (Figure 5E, 5F) among the CAR^+^ T cells derived from the PBMC donors. CD8^+^ T cells comprised the major T cell subtype population (Figure 5D), despite the known higher prevalence of CD4^+^ T cells in human PBMCs. To analyze this phenomenon, we carried out a time-course experiment to determine CD8/CD4 ratios before and up to 14 days following electroporation (Supplemental Figure 1). We categorized the time course into three stages: transient transfection (up to 5 days post-electroporation), stable transgene integration (after 5-7 days post-electroporation with undetectable transient CAR expression, Figure 4D), and cell expansion (10 and 14 days post-electroporation). As shown, the CD8 enrichment appeared to occur at the transfection (Supplemental Figure 1A and 1C) and the cell expansion stages (after seven days in culture), but not at the stable transgene integration stage (Supplemental Figure 1B, 1C and 1D). Furthermore, the major CAR^+^ T cell subset was T_SCM_ for both CD4 and CD8 populations (Figure 5E, 5F).

### *Quantum pBac*™ (*qPB*) system produces functional CAR^+^ T cells that kill target cells *in vitro*

We next sought to determine whether CAR-T cells generated using the *qPB* system (*qPB* donor vector with *qPBase v2* helper plasmid) are functional *in vitro*. We conducted *in vitro* cytotoxicity assays by co-incubating antigen-expressing target cells with either non-transfected control Pan-T cells or CD20/CD19 dual-targeting CAR-T cells (Figure 6).

**Figure 6.**
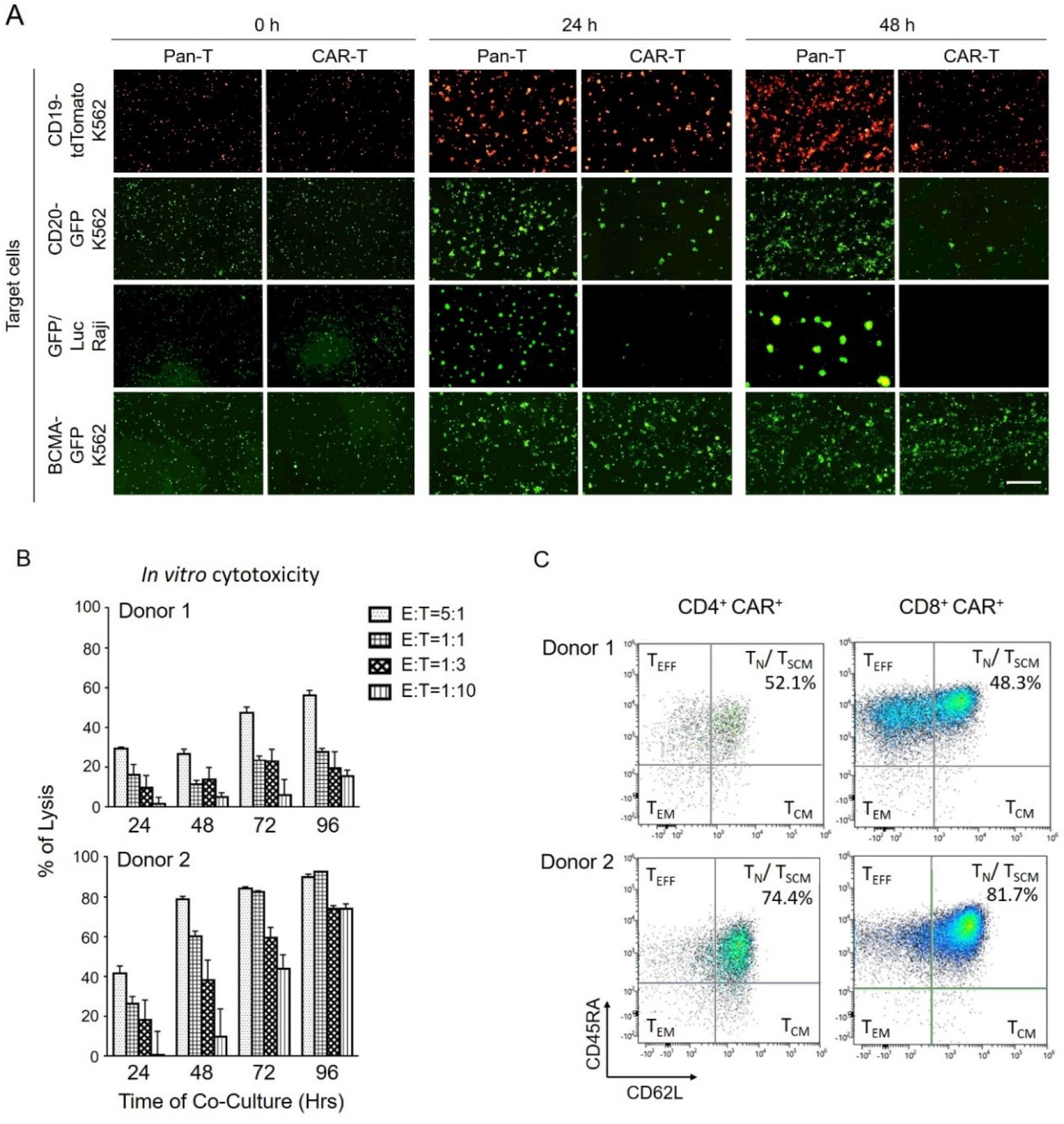
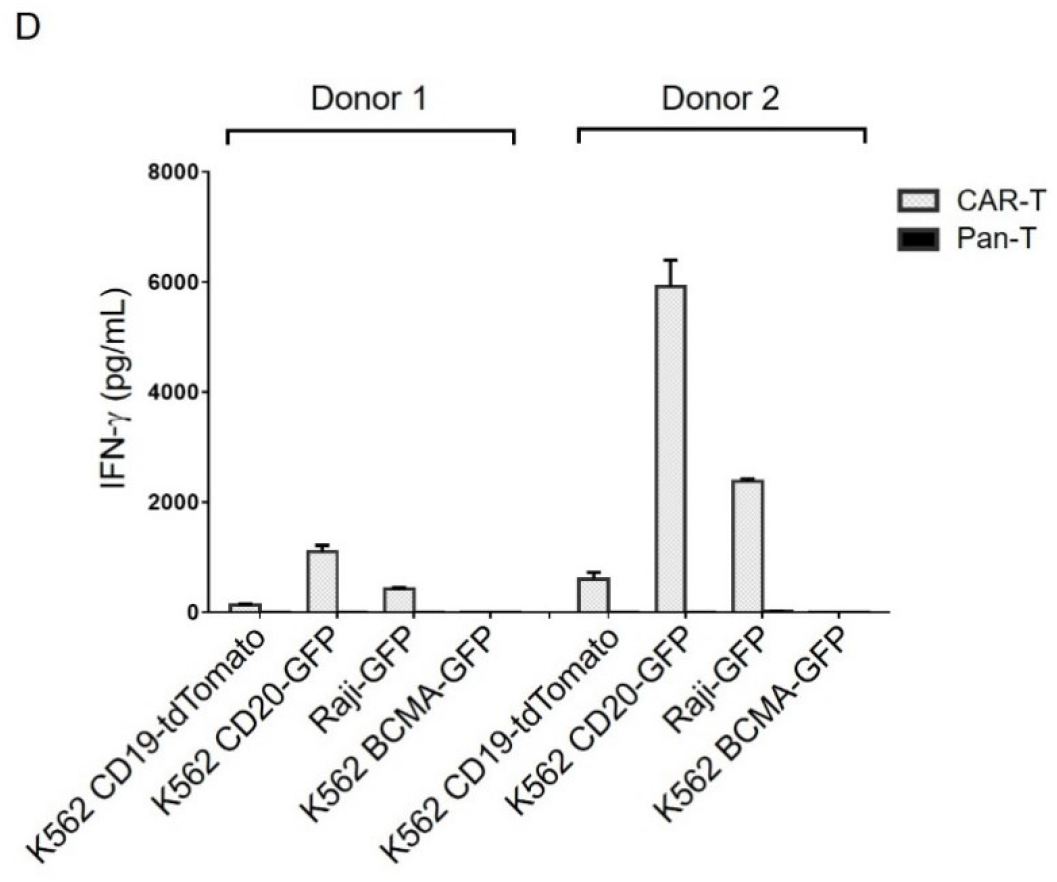
*In vitro* functional characterization of CAR-T cells produced using the *Quantum pBac*™ (*qPB*) system. *In vitro* cytotoxicity of Pan-T and/or CAR-T cells derived from healthy donor(s) (A) against the indicated target cells, and (B) against Raji cells at the indicated Effector: Target (E: T). (C) Representative flow cytometry data show the distributions of T cell differentiation subsets T_N_/T_SCM_, T_CM_, T_EM_ and T_EFF_ in CD4^+^ and CD8^+^ subtypes for donor 1- and donor 2-derived CAR^+^ T cells. (D) IFN-g secretion by donor 1 and donor 2 CAR-T cells following antigen stimulation. Pan-T cells (non-gene modified cells) served as a control. Data shown in (B and D) represent mean ± SD. N = 3 (triplicate). Bar in (A) represents 500 mm.

As shown in Figure 6A, compared to pan-T cells, CD20/CD19 dual-targeting CAR-T cells eradicated more CD19^+^CD20^+^ Raji cells (third row) and K562 target cells engineered to express either CD19 (first row) or CD20 (second row). On the other hand, CD20/CD19 dual-targeting CAR-T cells failed to eradicate BCMA (irrelevant antigen)-expressing control K562 cells (fourth row). These results demonstrated that the CAR-T cells specifically target and kill both CD20- and CD19-expressing cells. The cytotoxic functions of CAR-T cells from two donors (donors 1 and 2) were further assessed at different E:T ratios against Raji cells. We observed dose- and time-dependent killing by CAR-T cells derived from both donors (Figure 6B). Compared with donor 1-derived CAR-T cells, donor 2 CAR-T cells were much more potent in killing Raji cells, which was consistent with the relatively higher level of IFN-γ detected in the 48-hour culture medium of donor 2 CAR-T cells (Figure 6D). Notably, 74% of Raji cells were killed by donor 2 CAR-T cells after a 96-h co-culture period even at a E:T ratio of 1:10 (Figure 6B). This finding is in contrast to the 15.5% killing of Raji cells by donor 1 CAR-T cells at the same E:T ratio. Together, these results suggest a higher level of persistence for CAR-T cells derived from donor 2 as compared with those from donor 1. Supporting this notion, the percentages of CAR^+^ T_SCM_ cells were higher among CAR-T cells from donor 2 (74.4% and 81.7% for CD4^+^ and CD8^+^ cells, respectively) than those from donor 1 (52.1% and 48.3% for CD4^+^ and CD8^+^ cells, respectively; Figure 6C).

### Effective tumor clearance by *Quantum pBac*™ (*qPB*)-generated CAR-T cells in Raji-bearing immunodeficient mice

Next, we tested the *in vivo* anti-tumor potency of donor 1 and donor 2 CAR-T cells in Raji-bearing immunodeficient mice (Figure 7A). Similar to the *in vitro* cytotoxicity results, injections of low, medium and high doses of donor 1 CAR-T cells for five days killed Raji tumor cells in a dose-dependent fashion, and the tumors were completely eradicated by Days 5 and 9 in mice injected respectively with high and medium doses of donor 1 CAR-T cells (Figure 7B).

**Figure 7.**
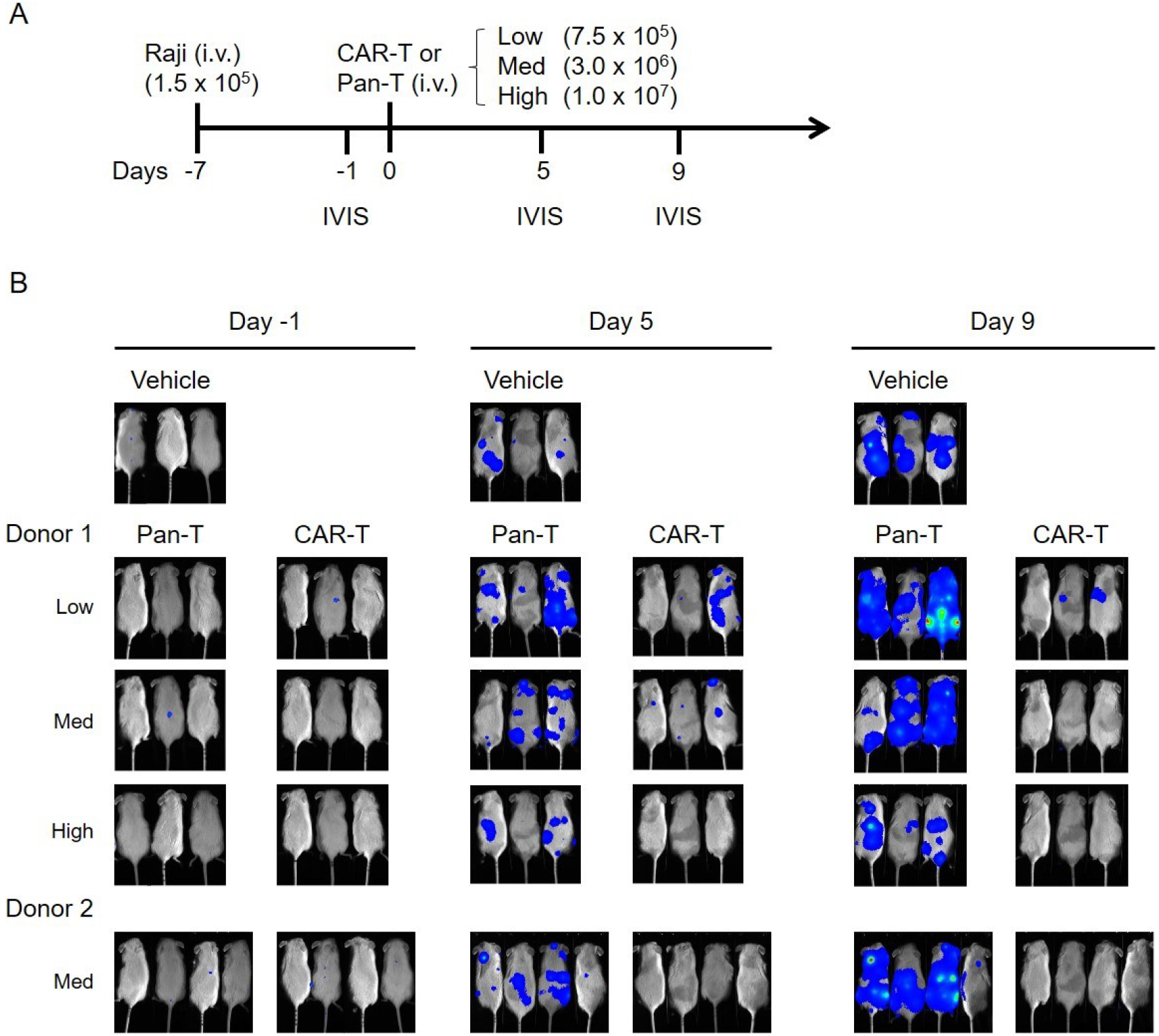
*In vivo* functional characterization of CAR-T cells produced using the *Quantum pBac*™ (*qPB*) system in Raji-bearing immunodeficient mice. (A) Schematic depicts the design of the *in vivo* functional characterization experiment. (B) Bioluminescence imaging (IVIS) revealed the extent of tumor cell persistence 1 day prior to, and 5 and 9 days after CAR-T or Pan-T cell injection. N = 3 or 4 mice/group.

Consistent with these findings, a medium dose of donor 2 CAR-T cells also eradicated Raji tumor cells from the Raji-bearing mice. Notably, the Raji tumor cell killing by donor 2 CAR-T cells appeared to have occurred at an earlier time point (at Day 5 following CAR-T cell injection) than that induced by donor 1 CAR-T cells (at Day 9 following CAR-T cell injection) (Figure 7B), in agreement with the high *in vitro* cytotoxicity induced by donor 2 CAR-T cells.

## Discussion

*Sleeping Beauty* and *piggyBac* are two DNA transposons that have been explored in recent clinical studies on gene and cell therapy^14,30^. In addition to the multiple advantages over viral vectors mentioned earlier, the *piggyBac* transposon system has at least four other major benefits. (1) It has a large cargo capacity (> 100 kb)^31^. (2) There is a low frequency of footprint-induced mutations caused by integrant remobilization^32–34^. (3) *piggyBac* is perhaps the most active transposon system in human cells^35^. (4) It is the most flexible transposon system in terms of retained activity upon molecular engineering of the transposase; this aspect may be crucial for applications that require modifications of the transposase to direct site-specific *piggyBac* genomic integration. These unique features also make *piggyBac* superior to *Sleeping Beauty* as a gene therapy vector. Nevertheless, *Sleeping Beauty* is thought to be less genotoxic than *piggyBac* for two main reasons: First, *piggyBac*-like terminal repeat elements are prevalent in the human genome^36^. Second, unlike the far more random genome integration profile of *Sleeping Beauty*, the genome integration profile of *piggyBac* is associated with euchromatin and largely excluded from heterochromatin^37,38^. The relatively high likelihood of *piggyBac* integration proximate to or within genes raises safety concerns, since enhancer elements of retroviral vector long terminal repeats (LTR) have previously been implicated as triggers of insertional mutagenesis in SCID-X1 patients^39^.

The first above mentioned safety concern of *piggyBac* system was thoroughly evaluated in a previous study,^40^ which demonstrated that the expression of transposase did not result in mobilization of endogenous *piggyBac*-like sequences from the human genome^40^. The authors also did not find a significant increase in double-stranded DNA breaks or a selective growth advantage in *piggyBac*-harboring cells within long-term cultures of primary human cells that were modified with eGFP-transposons^40^. To address the second above mentioned concern (related to the potential risk of insertional mutagenesis resulting from integrant enhancer activity), we determined whether enhancer activity was present in the TRDs and/or the TIRs. We discovered that the IDS region of 3’ TRD contained significant enhancer activity (Figure 1). By removing the IDS, our micro-*piggyBac* is expected to exhibit significantly lower levels of enhancer activity, which in turn may improve the safety profile of the donor vector. However, it has been well documented that both TIRs and other sequences contained within the TRDs are crucial for efficient integration of *piggyBac* transposon into the host genome. Several attempts to decrease the potential genotoxicity by reducing the size of the required TRDs to approximately 100 base pairs of TIRs have resulted in significant losses in transposition efficiency^27,41–43^.

In contrast to these findings, our previous study^25^ and the present study both demonstrated that by deleting IDS from both ends and leaving only TIRs (107 bp total in size), the transposition efficiency of *piggyBac* was in fact increased in several instances (Figure 2 and Figure 3).^25^

Importantly, a significant increase in transposition efficiency was observed in all cell types in which cells were co-transfected with a helper plasmid expressing *qPBase v2* when compared to those transfected with plasmid expressing wild type *PBase*. This finding suggests that *qPBase v2* is a robust and cell type-independent *piggyBac* transposase that can exhibit high transposition activity irrespective of the type and configuration of the *piggyBac* donor vector. We also demonstrated that *qPBase v2* mediated significantly higher transposition activities than *hyPBase* (1.8-fold increase in percentage of CAR^+^ cells on day 10 over *hyPBase*; Figure 4D), while their CAR gene copy numbers did not significantly differ. Our previous study has shown in a HEK293 genome-wide analysis of *qPBase v2*’s integration sites that *qPBase v2* displayed a much more random integration profile with no detectable hot spots, and has a lower preference for CpG islands and cancer-related genes than *qPBase v1* or wild-type *PBase*.^26^ Therefore, *qPBase v2*’s enhanced transposition activities over *hyPBase* may be due to an increased preference of *qPBase v2* for regions of the chromosome which promotes transgene expression. With regards to the potential reason why cells produced using *qPBase v2* expanded more than those produced using *hyPBase*, both the above mentioned significantly higher transposition activities of *qPBase v2* as well as the higher percentages of both CD4 and CD8 CAR^+^ T_SCM_ (Figure 4D) likely play a role. It is noteworthy that while the CAR copy numbers in our comparison experiments seemed relatively high, we have determined in a separate study related to manufacturing CAR-T cells for clinical applications that the total DNA electroporated is positively-associated with CAR copy number in CAR-T cells. We have subsequently optimized the donor:helper ratios to maximize generation of CAR^+^ T cells while maintaining acceptable CAR copy numbers.^44^ Collectively, the observations of this study suggest that micro-*piggyBac* in conjunction with *qPBase v2* comprises a superior *piggyBac* system that outperforms other *piggyBac* systems comprising *hyPBase*.

In addition to the impacts of TRDs/TIRs and transposase on the performance of the *piggyBac* system, the method of introducing these components into host cells can also greatly influence the transposition efficiency, potential safety profile, and transgene stability of the *piggyBac* system. To ensure co-transfection of both the transgene delivery cassette and the transposase gene, a single plasmid transposon system was previously created wherein TRDs were placed within the delivery cassette, and the transposase gene was located in the helper part of the same plasmid^27^. However, such an arrangement had a few disadvantages, including: (1) a reduced rate of gene delivery due to an increase in size of DNA, a phenomenon particularly seen in electroporation-based gene delivery, (2) a high degree of plasmid backbone DNA integration when using transposon plasmids^27^, and (3) a high rate of transposase gene integration into the host genome^45^. For these reasons, we opted to utilize a two-component system, consisting of donor vector and helper plasmid.

The use of a donor vector in plasmid form may have several undesirable qualities for clinical translation, including the presence of bacterial genetic elements and antibiotic-resistance genes. These features are undesirable since they pose a significant safety risk if the plasmid backbone becomes transposed into the host’s genome. Additionally, unmethylated CpG motifs that are highly enriched in the bacterial backbone of plasmids have been shown to trigger strong inflammatory responses in host cells through toll-like receptor-9 associated mechanisms. The sequences may also induce transgene silencing, presumably via cellular immune responses^46^ and/or interferon secretion^47^.

To reduce the chance that such events may occur, a new *piggyBac* system was established by incorporating *piggyBac* transposon into a doggybone™ DNA (dbDNA) vector, which is a linear, covalently closed, minimal DNA vector produced enzymatically *in vitro*^48^. A recent report has demonstrated that this *piggyBac*-incorporated dbDNA can be used to generate stable CD19-targeting CAR-T cells with an efficiency similar to that of its plasmid counterpart; the efficiencies were comparable when a minimum of approximately 470 bp of additional randomly selected DNA flanking the transposon was included. However, due to its linear configuration, dbDNA is expected to be less efficient for electroporation-based gene delivery as compared to its supercoiled circular counterpart. In this study, we set out to increase *piggyBac*’s cargo capacity by using Mark Kay minicircle technology to generate the smallest known *piggyBac* transposon donor vector with TIRs (107 bp) and only 88 bp of the backbone sequence flanking the TIRs. The small backbone sequence in combination with the truncated TRDs (i.e., TIRs) also resulted in a marked decrease (more than 2 kb) in the size of the donor vector. The smaller size donor vector in turn reduced DNA toxicity (i.e., electroporation-induced cell damage) and enhanced the transfection rate, which respectively improved cell expansion and transposition efficiency. This increase in transposition efficiency was further amplified when the donor vector carried a larger transgene (4.0 kb vs 5.34 kb). The end result of our modifications is that we now have a *piggyBac* system that is capable of successfully delivering and transposing larger size transgenes compared to *hyPB*. We demonstrated that use of this transposon donor vector in conjunction with a highly active *qPBase v2*, collectively designated as *Quantum pBac*™ (*qPB*), represents a superior *piggyBac* system that is minimalistic, highly efficient, and potentially safe (Figure 3).

Currently, the failure rate for producing sufficient number of CAR^+^ T cells for therapy is quite high, and this high rate may be traced back to the scarcity of T cells in some patients. Moreover, the T cells that remain in these patients are often exhausted and senesced, making the cells more difficult to expand to reach clinical quantity and quality criteria. In this study, we have addressed this obstacle with our *qPB* system utilizing *qPBase v2*. This system can be utilized to effectively generate CD19/CD20 dual-targeting CAR-T cells with a 1.8-fold increase in the percentage of CAR^+^ cells on day 10, and a 2-fold increase in the 10-day expansion capacity, compared to *hyPBase* (Figure 4). Together, these increases in CAR-T cell performance result in a 3.6-fold increase in the final yield of CAR-T cells. We showed that this difference in yield is likely due to an enhanced transposition efficiency achievable by the *qPB* system (Figure 3), and the increased yield should facilitate the rapid generation of CAR^+^ cells at a clinical scale. Likewise, the significantly higher cytolytic activities of CAR-T cells produced using *qPBase v2* compared with those produced using *hyPBase* (Figure 4G) likely reflect the significantly higher percentage of CAR^+^ cells in the cells produced by *qPBase v2* than by *hyPBase* (Figure 4D) and greater proportion of T_SCM_ CAR^+^ T cells (Figure 4E). This advance in technology may prevent the need to extend CAR-T cell production time; such extensions may be required when using other CAR-T production systems and may increase the risks of T cell exhaustion and senescence that compromise CAR-T quality. The higher proportion of CAR^+^ T cells that are produced by using the *qPB* system should also allow for injection/administration of fewer total cells, which may reduce the adverse effects of CAR-T therapy. We have also shown that as the transgene size increased, the percentage of CAR^+^ cells decreased.^49^ Since the difference in transposition efficiencies between plasmid backbone-containing and plasmid backbone-lacking donor vectors became more pronounced as transgene size increased (4.0 kb in Figure 2D vs 5.34 kb in Figure 3C), we should be able to improve therapy results by reducing the size of the donor vector to increase its capacity for therapeutic transgene payload. Therefore, use of the *qPB* system, which is capable of carrying a larger transgene cargo due to its reduced size, should enable the production of efficacious CAR-T cells harboring sizable transgene(s) at a clinical scale.

It is noteworthy that there was a high representation of CD8^+^ T cells in CAR^+^ T cells produced using the *qPB* system (Figure 5D), despite the known high prevalence of CD4^+^ T cells in human PBMCs. As shown in Supplemental Figure S1A, the CD8/CD4 ratio initially increased within one day post-electroporation, suggesting that CD4 T cells may be more sensitive to electroporation-associated damages. Subsequently, the further increase in CD8/CD4 ratio was positively associated with a marked increase in cell expansion (see Figure S1C and S1B, respectively), suggesting that the CD8 T cells were preferentially expanded compared to the CD4 T cells. The fact that there was no change in CD8/CD4 ratio at 5 and 7 days post-electroporation (when only transposed and not transfected transgenes could be detected) also suggests that the pronounced enrichment of CD8 T cells was not due to preferential transposition of CD8 T cells. As demonstrated in another study, this preferential enrichment of CD8 T cells can serve to complement the more difficult-to-enrich patient CD8 T cells and contribute to a balanced CD8/CD4 ratio (approximately 1.0) in CAR-T cells produced from patients with B-cell malignancies.^44^

Additionally, CAR-T cell expansion and persistence have emerged as key efficacy determinants in cancer patients, and both are positively correlated with the proportion of T_SCM_ cells in the final CAR-T cell product^50,51^. In line with these findings, we have demonstrated that our *qPB*-derived CAR-T cells, which are enriched in T_SCM_ cells (especially among CD8^+^ CAR^+^ T cells), exhibited potent anti-tumor effects both *in vitro* and *in vivo*.

Another strategy to improve CAR-T therapy is to minimize genotoxicity caused by transposase-induced integrant remobilization. To evaluate this potential approach, transposase mRNA has been examined in transposon-based CAR-T clinical trials^52^. However, compared to transposase DNA, the mRNA is generally less efficient, more costly (for GMP production), and less stable under storage conditions. Given the low frequency of footprint-induced mutation by *piggyBac* (<5%) and the fact that only a minimal proportion (<0.4%) of CAR^+^ T cells expressed detectable levels of *qPBase* (Figure 5B), transposase in a plasmid form should be sufficiently safe when applied to highly proliferative *ex vivo*-engineered cells. Additionally, our recent genome-wide integration profiling of *qPB* in CAR-T products derived from two distinct donors further support the notion that it is safe to use in T cell engineering^49^.

In summary, given its larger payload capacity compared to currently available *piggyBac* systems and high level of efficiency, *qPB* is likely the most suitable system for development of next generation virus-free gene and cell therapies, especially for multiplex CAR-T therapy. As we have demonstrated in this study, the *qPB* system exhibits greater transposition activity, causes less damage to cells, induces a higher percentage of T_SCM_ and CAR^+^ populations, and expands more of the engineered cells than previous systems. These attributes lead to an overall enhanced ability of the *qPB* system to produce highly CAR^+^ T_SCM_ cells harboring larger payloads, which should contribute to the success of *piggyBac*-based multiplex CAR-T gene therapy. The success of *qPB* in CAR-T cell therapy suggests that the technology may also be suitable for use in other gene therapy applications as well.

## Materials and Methods

### Human T cell samples from healthy donors

Blood samples from adult healthy donors were obtained from Chang Gung Memorial Hospital (Linkou, Taiwan). The acquisition of these samples was approved by the Institution Review Board at Chang Gung Medical Foundation (IRB No. 201900578A3).

### Vector constructs

Plasmids were constructed following protocols described previously^53^. Minicircle DNAs were purchased from Aldevron (Fargo, ND). All of the PCR products or junctions of constructs (wherever sequences were ligated) were confirmed by sequencing.

### Vector construction

#### pGL3-miniP (Construct a; Figure 1A)

pGL3-basic was digested with SacI and HindIII. The CMV mini-promoter with SacI and HindIII sequences on its 5’ and 3’ ends, respectively, was synthesized, double-digested with SacI and HindIII, and ligated to the SacI-HindIII vector fragment of pGL3-basic to make construct **a** as shown in Figure 1A. **Constructs b-e (Figure 1A)**

pGL3-miniP was digested with KpnI and XhoI. The 5’TIR or 3’TIR of micro-*piggyBac* or 5’TRD or 3’TRD of mini-*piggyBac* with KpnI and XhoI sequences added on either end was synthesized, double-digested with KpnI and XhoI, and ligated to the KpnI-XhoI vector fragment of pGL3-miniP, respectively, to make the set of constructs (**b**-**e**) for evaluation of enhancer activity of mini-*piggyBac* and micro-*piggyBac* TRDs and TIRs, respectively.

#### Donor vectors (Figure 2A)

“Mini-*piggyBac*-Long” was the same construct as the donor of *piggyBac* system published previously ^19^. “Micro-*piggyBac*-Short” was the plasmid named “*pPB*-cassette short” in a previous publication^25^. “Micro-*piggyBac*-Long” was constructed by replacing the backbone of “micro-*piggyBac*-Short” with that of “mini-*piggyBac*-Long” via PCR-based cloning.

#### Helper plasmid (Figure 2A)

The construction of helper plasmid was described previously^26^.

#### Parental *qPB* donor vector (Figure 3A)

The parental plasmid contained the following components (obtained by DNA synthesis) arranged in a 5’ to 3’ order: Kanamycin resistance gene, origin of replication (pMB1), 32 I-SceI sites, attB site (PhiC31), 5’TIR of micro-*piggyBac*, multiple cloning site (MSC), 3’TIR of micro-*piggyBac*, and attP site (PhiC31).

#### pQ-tdTomato-IRES-hygro (Figure 3A)

The DNA for a cassette with the CAG promoter driving a bi-cistronic transcript containing tdTomato gene, internal ribosome entry site (IRES) of Foot-and-mouth disease virus (FMDV), and hygromycin resistance gene was synthesized and cloned into the AseI and EcoRV sites of the parental *qPB* vector to make pQ-tdTomato-IRES-hygro.

#### pQ-CD20CD19CAR (Figure 4A)

The dual-targeting tandem CD20/CD19 CAR-containing pQ-CD20CD19CAR vector encoded a second-generation CAR composed of an elongation factor 1-alpha (EF1α) promoter that drives an extracellular domain derived from the single chain variable fragments (scFv) of monoclonal antibodies directed against the CD19 and CD20 antigens, respectively. The sequence is further linked to the CD3z chain of the TCR complex by means of a CD8 hinge and transmembrane domains, together with the 4-1BB co-stimulatory domain. Following synthesis, the expression cassette was cloned into the MCS of parental *qPB* vector to make pQ-CD20CD19CAR.

#### Minicircle DNA

The minicircle DNAs, Q-tdTomato-IRES-hygro (Figure 3A) and Q-CD20CD19CAR (Figure 4A), were manufactured by Aldevron from parental plasmids, pQ-tdTomato-IRES-hygro and pQ-CD2019CAR, respectively.

#### qPBase

To comply with the FDA regulations, the ampicillin gene in the helper plasmid expressing *qPBase v2* was replaced by a kanamycin resistance gene sequence to make *qPBase*. The *qPBase* component was combined with *qPB* donor vector (minicircle DNA) to form the *qPB* system.

#### Enhancer assay

The control, pPL-TK (Renilla Luciferase), was co-transfected with the specified firefly luciferase constructs (i.e., pGL3-miniP, pGL3-miniP-microL, pGL3-miniP-microR, pGL3-miniP-miniR, pGL3-miniP-miniL, mini-*piggyBac* Long, micro-*piggyBac* Short, or micro-*piggyBac* Long) by either FuGENE (HEK293) or nucleofection (Sf9 cells, and Jurkat T cells). In some experiments, constructs were first linearized using XmnI or BglI. Forty-eight hours after transfection, cells were harvested and subjected to the Dual-Luciferase assay (Promega) according to the manufacturer’s instructions.

#### Transposition assay

HEK293 cells (1 × 10^5^) were transfected with 200-334 ng of donor vector carrying a hygromycin resistance gene and 200-282 ng of helper plasmid in MEM medium (GeneDireX), 10% FBS (Corning) utilizing the X-tremeGENE™ HP DNA transfection reagent (Merck). Transfected cells were transferred to 100-mm plates and cultured under hygromycin (100 μg/ml) selection pressure for 14 days. Cells were then harvested and fixed with 4% paraformaldehyde. Fixed cells were stained with 0.2% methylene blue and cell colonies were counted.

Jurkat or primary T cells (2 × 10^5^) were electroporated with 400-668 ng of donor vector carrying a hygromycin resistance gene, which in some experiments was also linked to a tdTomato gene, and 400-517 ng of helper plasmid in OpTmizer medium supplemented with *Quantum Booster*™ (GenomeFrontier). Electroporation was carried out using a 4D-Nucleofector™ (Lonza) in combination with the *Quantum Nufect*™ Kit (GenomeFrontier) according to the manufacturer’s instructions. Electroporated cells were transferred to 96-well plates and cultured under hygromycin (1 mg/ml) selection pressure for 14 days. Cells were then harvested and stained with AO/PI. Live cell numbers were determined using Celigo image cytometry (Nexcelom). Transposition efficiency was expressed as number of either hygromycin-resistant colonies or live cells.

#### Generation and expansion of CAR-T cells

Peripheral blood mononuclear cells (PBMCs) were isolated from blood samples of healthy donors by Ficoll-Hypaque gradient separation. CD3^+^ T cells were isolated from PBMCs using EasySep™ Human T Cell Isolation Kit (StemCell Technologies) according to the manufacturer’s instructions. T cells were activated by co-incubation with anti-CD3/anti-CD28 antibody conjugated Dynabeads™ (Gibco, Catalogue No. 11141D) in X-VIVO 15 medium (Lonza) for two days at a bead to cells ratio of 3:1. Following the removal of Dynabeads™, activated T cells were harvested and frozen or utilized in experiments. Electroporation of activated T cells was carried out using a Nucleofector™ 2b Device (Lonza) in combination with the *Quantum Nufect*™ Kit (GenomeFrontier) according to the manufacturer’s instructions. Cells were electroporated with the combination of CD20/CD19 CAR donor vector, and pCMV-hyPBase or pCMV-qPBase v2 helper vector. Cells were cultured and expanded for 10 or 14 days in OpTmizer medium (Thermo Fisher Scientific) supplemented with 50 IU of IL-2 (PeproTech) and 10% FBS. Thereafter, the cells were harvested for experiments. γ-irradiated aAPCs were added on Day 3 to the T cell expansion cultures at an aAPC:T cell ratio of 1:1.

#### Evaluation of CAR-T cells performance

CAR expression on T cells was detected by flow cytometry following staining of cells at 4°C for 30 min with F(ab’)_2_ fragment-specific, biotin-conjugated goat anti-mouse antibodies (Jackson ImmunoResearch Laboratories) and R-phycoerythrin (PE)-conjugated streptavidin (Jackson ImmunoResearch

Laboratories). Similarly, cells were stained with the following antigen-specific fluorophore-conjugated antibodies: CD3-Pacific Blue, CD4-Alexa Flour 532 (Thermo Fisher Scientific), CD8-PE-Cy7, CD45RA-BV421, CD62L-PE-Cy5, or CD95-BV711 (Biolegend). In some experiments, cells were incubated with propidium iodide (PI, Thermo Fisher Scientific) and/or Acridine orange (AO, Nexcelom). PI-cells and T cell differentiation subsets were determined by flow cytometry based on CD45RA, CD62L and CD95 expression, as follows: T_N_ (CD45RA^+^CD62L^+^CD95^-^), T_SCM_ (CD45RA^+^CD62L^+^CD95^+^), T_CM_ (CD45RA^-^ CD62L^+^), T_EM_ (CD45RA^-^CD62L^-^), and T_EFF_ (CD45RA^+^CD62L^-^). *qPBase* expresses a GFP-transposase fusion protein, which was detected by flow cytometry for GFP^+^ cells. In some experiments, a hygromycin resistance gene was expressed with a tdTomato gene and hygromycin-resistant cells were identified by flow cytometry for tdTomato^+^ cells. Flow cytometric measurements and analyses were performed on a SA3800 Spectral Analyzer (Sony). Histograms and dot-plots were generated using GraphPad Prism software (GraphPad). Live cells were detected using a Celigo imaging cytometer (Nexcelom) as AO^+^, PI-cells.

#### *In vitro* cytotoxicity assay

Target-antigen-expressing cells were engineered according to a method described elsewhere^49^. CD19^+^ (K562 CD19-tdTomato), CD20^+^ (K562 CD20-GFP), CD19^+^CD20^+^ (Raji-GFP) or non-relevant K562 BCMA-GFP target cells were seeded at 5 × 10^3^ cells per well in 96-well culture plates (Corning), and control non-transfected Pan-T or CAR-T cells were added at an E:T ratio of 5:1, 1:1, 1:3 or 1:10. Cytotoxicity of CAR-T cells on target cells was then assessed as previously described^54^. Briefly, a Celigo imaging cytometer was used to measure the percentage of live target cells at 0, 24, 48, 72 and 96 h of co-culture. Cell aggregates were separated by pipetting prior to Celigo imaging. The percent of specific lysis for each sample was calculated using the formula: [1-(live fluorescent cell count in the presence of target cells and CAR-T cells / live fluorescent cell count in the presence of target cells only)] x 100.

#### *In vitro* cytokine release assay

Pan-T control cells or CAR-T cells produced from two healthy donors were thawed and added to cultures containing CD19^+^ (K562 CD19-tdTomato), CD20^+^ (K562 CD20-GFP), CD19^+^CD20^+^ (Raji-GFP) or non-relevant K562 BCMA-GFP tumor target cells. The cells were added at an effector:target (E:T) ratio of 10:1 in OpTmizer medium supplemented with 50 IU of IL-2 and 10% FBS. After 48 h of co-culture, supernatant was collected and IFN-γ level in the culture supernatant was measured by an enzyme-linked immunosorbent assay (Thermo Fisher), according to the manufacturer’s instructions.

#### Mouse xenograft model

*In vivo* studies on a mouse xenograft model were conducted at the Development Center for Biotechnology, Taiwan, using animal protocols and procedures ethically reviewed and approved by the Taiwan Mouse Clinic IACUC (2020-R501-035). Briefly, 8-week-old female ASID (NOD.Cg-Prkdc^scid^Il2rg^tm1Wjl^/YckNarl) mice (National Laboratory Animal Center, Taiwan) were intravenously (i.v.) injected with 1.5 × 10^5^ Raji-Luc/GFP tumor cells. One week after Raji-Luc/GFP tumor cell injection, mice were injected with Low (7.5 × 10^5^), Med (3 × 10^6^), or High (1.0 × 10^7^) doses of CAR-T cells or control Pan-T cells.

Luminescence signals from Raji-Luc/GFP tumor cells were monitored using the Xenogen-IVIS Imaging System (Caliper Life Sciences).

### Statistical analysis

Statistical analyses of differences between two groups were carried out using Student’s t-test (two-tail), and differences among three or more groups were analyzed by one-way ANOVA with Tukey’s multiple comparison test. The analyses were performed using GraphPad Prism software (GraphPad Software), and statistical significance was reported as * p < 0.05, ** p < 0.01, and *** p < 0.001. Differences were considered to be statistically significant when p < 0.05.

## Acknowledgments

The authors thank Ms. Lu-Chun Chen for her assistance throughout the IRB preparation and approval process. The authors also thank Dr. Pei-Yi Tsai for assistance with the animal experiments. This study was funded by GenomeFrontier Therapeutics, Inc.

## Author Contributions

S.C.-Y.W. designed research. Y.-C.C. (Yi-Chun Chen), P.-N.W., Y.-S.Y., Y.-C.C. (Ying-Chun Chen), I.-C.C. performed research. W.-K.H., Y.-C.C. (Yi-Chun Chen), J.C.H., K.-L.K.W., and S.C.-Y.W. analyzed data. J.C.H., P.S.C. and S.C.-Y.W. wrote the paper.

## Data Availability Statement

The authors confirm that the data supporting the findings of this study are available within the article and/or its supplementary materials.

