## Supplementary Materials for "*Quantum pBac*: An effective, high-capacity *piggyBac*-based gene integration vector system for unlocking gene therapy potential"

**This supplementary information file contains:**

(1) Supplementary Figure Legend; and

(2) Figure S1

**Supplementary Figure Legend**

**Supplementary Figure S1. Changes in CD8:CD4 ratios during CAR-T production.**

$CD3^+$ T cells were isolated and then activated for two days using Dynabeads. Activated T cells were nucleofected with the CD20/CD19 CAR *qPB* donor vector and the indicated helper plasmid. (A, C, and D) The CD8:CD4 ratios and (B) cell growth were determined on the indicated day of culture. Results are shown as mean ± SD. * $p < 0.05$, ** $p < 0.01$, *** $p < 0.001$. N = 3 (triplicate).

**Figure S1**

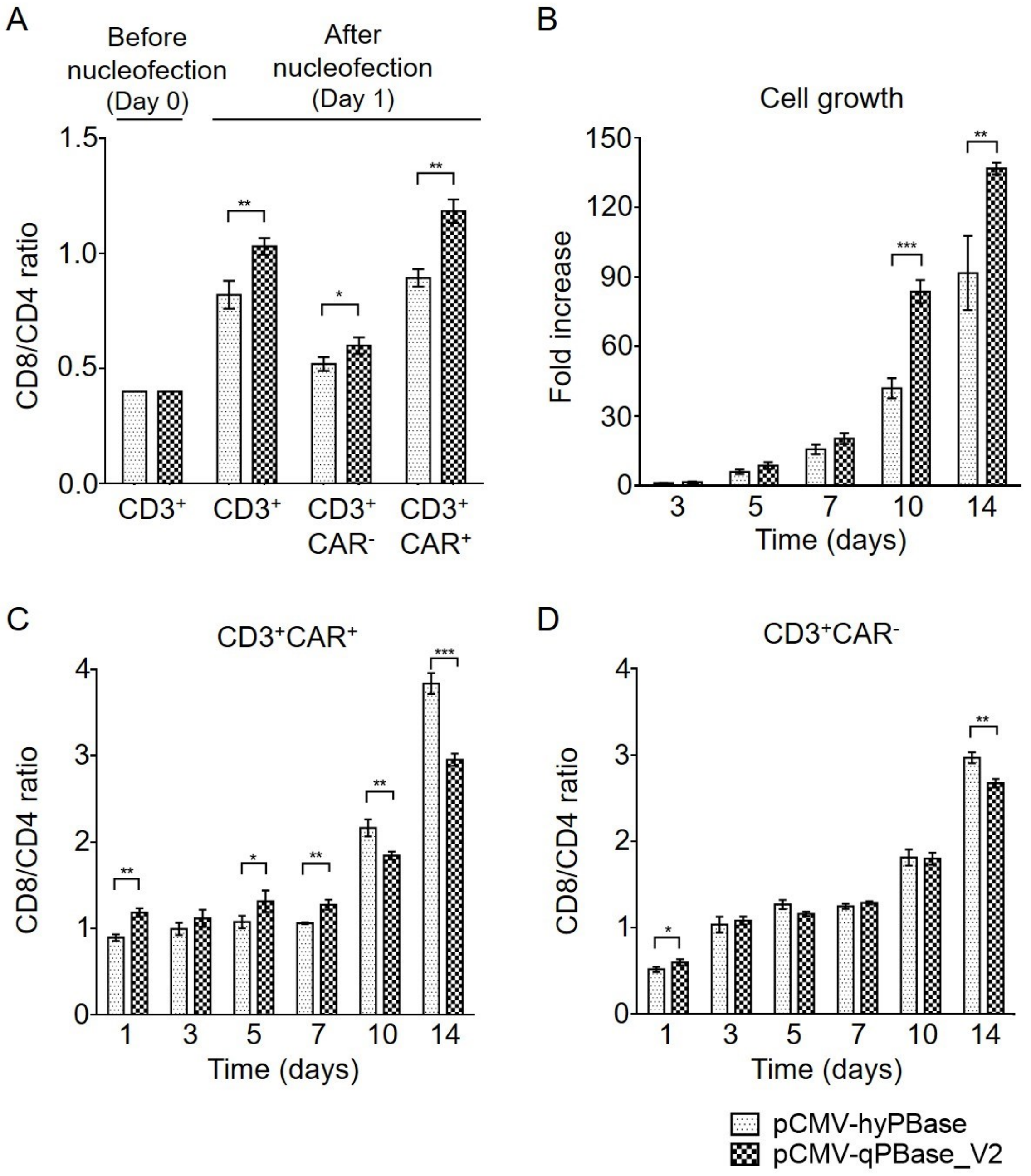

A
Before nucleofection (Day 0)
After nucleofection (Day 1)
CD8/CD4 ratio
1.5
1.0
0.5
0.0
CD3+
CD3+
CD3+ CAR-
CD3+ CAR+
B
Cell growth
Fold increase
150
120
90
60
30
0
3
5
7
10
14
Time (days)
C
CD3+CAR+
CD8/CD4 ratio
Time (days)
D
CD3+CAR-
CD8/CD4 ratio
Time (days)
pCMV-hyPBase
pCMV-qPBase_V2
